# Delineating Mutation Bias and Selection during Plant Development

**DOI:** 10.1101/2025.11.01.686053

**Authors:** J Grey Monroe, Mariele Lensink, Vianney Ahn, Matthew W. Davis, Satoyo Oya, Kehan Zhao

## Abstract

Plants accumulate germline mutations through meristematic lineages where somatic purifying selection could remove deleterious coding mutations and produce perceived mutation biases, like lower genic mutation rates. Here we describe a quantitative framework that uses mutation spectra and genome architecture to separate missing coding mutations from true genic hypomutation for selection-corrected estimates of mutation biases. In simulations of plant meristems subject to selection and mutation bias, only extremely strong dominant mutations produced gene-body patterns in the same direction as empirical estimates of hypomutation. However, after accounting for this selection, we can recover the true underlying mutation rate variation. We apply this to re-analysis of *Arabidopsis* mutation-accumulation datasets. After selection, we see ∼50% lower mutation rates in gene bodies and an additional ∼15–30% reduction in broadly expressed genes, consistent with mutation rate bias. This framework provides a quantitative method to compare mutation rates across the genome while explicitly controlling for selection on coding sequence mutations.

## Introduction

Mutation rates govern the arrival of new genetic variation and are not uniform across genomic regions and contexts. Accurate measurements of mutation rate heterogeneity across organism genomes are crucial for making robust population genetic inferences and studying the mechanisms that shape intragenomic mutational landscapes. A challenge in studying mutation rates is distinguishing the effects of mutation biases from selection – whether at the organism level or among cells, as in the case of competitive somatic mosaicism during development.

Mutation accumulation experiments in the model plant *Arabidopsis thaliana* (Arabidopsis, hereafter) measure mutation rates after generations of imposing extreme population bottleneck events to reduce the efficacy of selection. In plants, this is usually achieved by single seed descent, which reduces the effective population size to 1 at each generation, diminishing the efficacy of selection against all but lethal mutations. Despite these efforts, the possibility that observed mutation distributions are contaminated by the effects of cryptic selection, such as through competition among cells carrying different somatic mutations during development, is a challenge for mutation research (Majic & Payne, 2023). Moreover, in long-lived plants or mutation datasets not originally intended for accumulation studies, accounting for somatic selection is essential to accurately interpret observed mutational patterns (Y. Chen et al., 2024; Johannes, 2025b, 2025c; Schoen & Schultz, 2019).

Over the past decade, distinguishing genuine mutation rate variation from the effects of selection has been brought into sharp focus (B. Charlesworth & Jensen, 2022). We know now that DNA repair proteins, mediated by mechanisms like epigenome recruitment and transcription-coupled repair, can preferentially protect actively expressed genes from mutation (Aymard et al., 2014; Belfield et al., 2018; Fang et al., 2021; Huang et al., 2018; Li et al., 2013; Monroe et al., 2022; Moore et al., 2021; Quiroz et al., 2024; Sun et al., 2020; Supek & Lehner, 2015, 2017, 2019; Xiao et al., 2021). For example, transcription-coupled DNA repair in some tissues can cause an 80% reduction in the somatic mutation rate of the most highly expressed genes in humans (Moore et al., 2021; Nicholson et al., 2024). Exons also have consistently lower mutation rates in somatic tissue, due to elevated mismatch repair or nucleotide excision repair in humans (Frigola et al., 2017; Moore et al., 2021; Polak et al., 2014). In plants, reduced mutation rates of certain genomic regions have been repeatedly observed in mutation experiments (Davis et al., 2025; Goel et al., 2024; Meyer et al., 2025; Monroe et al., 2022; Ossowski et al., 2010; Perez-Roman et al., 2022; Quiroz et al., 2024). If these mechanisms were not well understood, their consequential genic hypomutation due to differential repair could be misperceived as the effects of selection, since genic regions are also the primary targets of purifying selection. Thus, mechanistic insights gained from mutation experiments can be uncertain if selection effects are not estimated and, if needed, accounted for in order to generate robust measures of mutation rate heterogeneity.

Selection can act both between generations and during development amongst somatic cells. The strength of purifying selection has been studied extensively in large somatic mutation datasets in humans, and largely reveals that purifying selection is weak or entirely absent during development, with the notable exception of hemizygous genome regions where the effects of recessive deleterious alleles are not masked by a complementary allele (Lawson et al., 2024; Martincorena et al., 2017). In inbred plants, like *Arabidopsis thaliana*, the strength of purifying selection is predicted to be especially weak due to the absence of hemizygosity (Abbott & Gomes, 1989; Platt et al., 2010). Nevertheless, considering selection both between generations and during development is important for a robust study of mutation rate heterogeneity (Majic & Payne, 2023).

Here, we outline a quantitative framework to estimate the relative effects of selection using non-synonymous and synonymous mutations to accurately infer biological mutation rate differences that underlie mutational cold spots in genic regions across organism genomes. We then apply this to test the conditions where observed genic hypomutation could be due to developmental selection. Finally, we apply this to a meta-analysis of mutation accumulation experiments in Arabidopsis and quantify the relative contribution of selection and mutation rate underlying gene body hypomutation.

## A framework to separate mutation heterogeneity from selection against non-synonymous mutations

### Quantifying selection against non-synonymous mutations

Non-synonymous mutations constitute the main targets of negatively selected sites in gene bodies (Haudry et al., 2013; Kimura, 1977; Miyata & Yasunaga, 1980; Z. Yang, 2007a). Therefore, to study the phenomenon of gene body hypomutation, one must consider the potential for selection on non-synonymous mutations to artificially deflate mutation rate measurements.

To evaluate whether observed gene-body mutation data are biased by purifying selection on nonsynonymous sites, we compare the observed nonsynonymous-to-synonymous count ratio (N_NS_^obs^/N_S_^obs^) with a neutral or codon/spectrum-based ratio R_NS/S_^(0)^. This allows us to distinguish (i) true genic hypomutation from (ii) apparent reductions caused by selective loss of nonsynonymous mutations.

#### Notation

μ_ref_ : mutation rate in the reference / intergenic region

μ_G_ : observed mutation rate in gene bodies

L_G_ : total length of genic sequence considered

P_CDS_ : fraction of gene bodies that is coding sequence (CDS)

R_NS/S_^(0)^ : neutral nonsynonymous-to-synonymous ratio. This ratio can be estimated empirically by determining the relative probability of NS versus S mutations arising in coding regions based on mutation spectra and codon use. Alternatively it can be measured directly if mutation rates are compared between genes, where the reference provides the neutral ratio.

N_NS_^obs^, N_S_^obs^, N_NC_^obs^ : observed counts of nonsynonymous, synonymous, and noncoding genic mutations

*E*μ_ref_[·] : expectation under the reference mutation rate model

*E*_S_[·] : expectation under neutrality, based on the observed number of synonymous mutations

#### Observed genic reduction

If gene-body mutation rates μ_G_ are reduced relative to the reference rate μ_ref_, we define the observed genic reduction factor as:

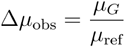

For example, Δμ_obs_ = 0.25 would indicate a 75% reduction.

#### Neutral composition of gene bodies

Let P_CDS_ be the CDS fraction of genes. Given a neutral nonsynonymous-to-synonymous ratio R_NS/S_^(0)^, the neutral expected proportion of nonsynonymous genic mutations is:

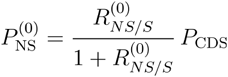

The corresponding expected proportions of synonymous and noncoding genic mutations are:

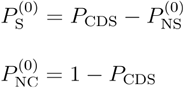

#### Reference / rate-based expected counts

Assuming genes mutate like the reference region (rate μ_ref_) and follow the composition above, the expected genic mutation counts under uniform mutation rates (no mutation bias) are:

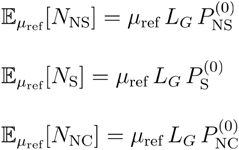

#### S-anchored expected nonsynonymous counts

The observed synonymous count provides a basis for an expected N_NS_ under neutrality (no selection) which is:

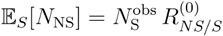

#### Testing for selection on nonsynonymous mutations

If the observed ratio is at least the neutral ratio,

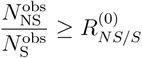

then selection on nonsynonymous mutations alone is unlikely to explain genic hypomutation.

If instead,

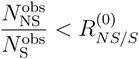

then nonsynonymous mutations are underrepresented and we can estimate how many have been removed.

#### Amount of NS removed and correction term

The number of missing nonsynonymous mutations is:

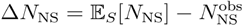

and the corresponding selection-correction term across the genic region is:

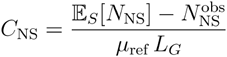

This is the number of NS missing due to selection (expected-from-S minus observed) per unit of total genic mutations expected under the reference rate.

#### Selection-corrected genic reduction

We then correct the observed reduction by adding back the selection term:

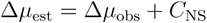

#### Component-wise hypomutation

To check whether mutation rates individual classes are also reduced, we compute:

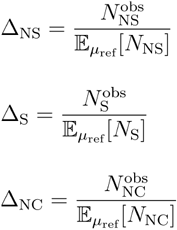

When Δ_S_ or Δ_NC_ are less than 1 and less than or equal to Δ_NS_, then a true, selection independent, genic mutation-rate reduction is further supported.

## Testing performance with simulations that model developmental selection and mutation bias in plant germline

### Simulation of germline

It was recently proposed that the genic hypomutation observed in *Arabidopsis thaliana* could be explained by developmental selection, specifically through competition between somatic cells that ultimately give rise to the germline *(Majic & Payne, 2023)*. However, this model did not consider the dominance or strength of selection on deleterious alleles, nor the specifics of plant development informed by meristem biology (Dai et al., 2024; Furner & Pumfrey, 1993; Huber et al., 2018; Lanfear, 2018). To study this hypothesis with more refined models, we constructed a forward population evolution simulation with SLiM (Haller & Messer, 2023) that corrects for previous limitations. In short, we simulated a population of clonally reproducing meristem cells, affected by varying degrees of purifying selection against genic mutations and true biological (i.e., DNA repair) hypomutation rate in genic regions (Table S1, **Figure 1a**). From this, and using the models discussed above, we aimed to test 1) under what conditions can developmental selection reduce genic mutations and 2) to what extent can correcting for purifying selection derive a more robust estimate of the true degree of genic hypomutation.

**Figure 1.**
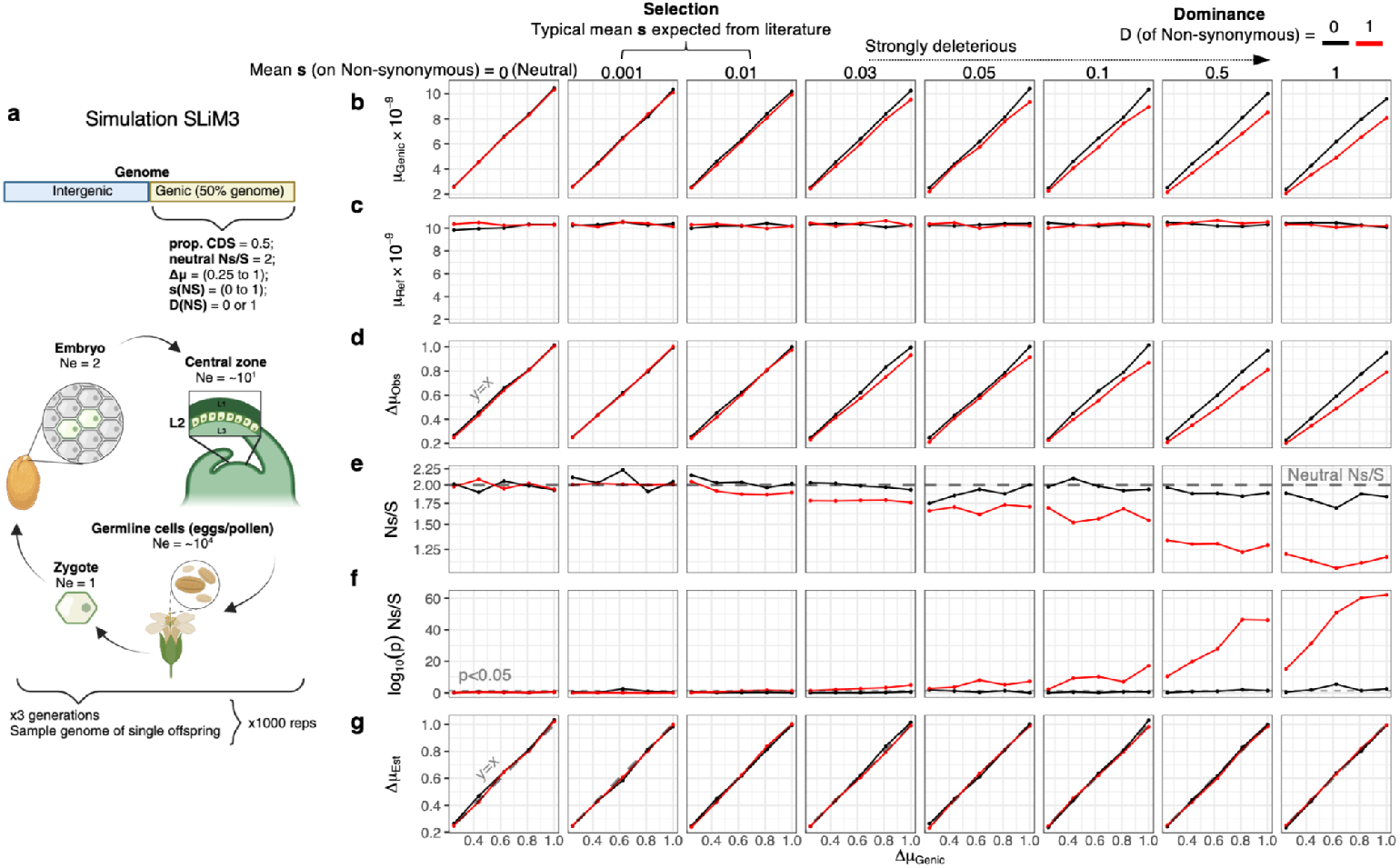
Simulation of selection and mutation rate bias in Arabidopsis mutation accumulation experiments. **a)** Overview of the simulation framework conducted using SLiM, modelled after the meristem development of the Arabidopsis from zygote to gamete, **b-g)** the results of these simulations, where the x-axis represents the true degree of mutation rate reduction (Δ μ) in gene bodies. The panels show the impact on measures of mutation bias and selection from various degrees of selection strength (s) and dominance (D) of *de novo* non-synonymous mutations: **b)** observed mutation rate (mutations per base pair per generation) in genic regions (scaled fo *A. thaliana* genome size) **c)** observed mutation rate in intergenic regions, **d)** the perceived reduction of mutation rate in gene bodies, with the dashed line indicating the true reduction, **e)** the observed non-synonymous to synonymous (NS/S) mutation ratio, with the dashed line representing the neutral expectation, **f)** results of chi-squared tests of NS/S ratio relative to the neutral expectation, indicating the significance of deviation from neutrality (p < 0.05, i.e. “*would we be able to detect selection*?’’) and **g)** the estimated reduction of mutation rate in gene regions after accounting for selection on coding mutations, comparing the estimated and actual mutation rate reductions.

In Arabidopsis, the number of genetically effective stem cells in embryos (cells that contribute to the germline) has been empirically determined to be 2 (Canales et al., 2002; Furner & Pumfrey, 1993; van den Berg et al., 1995). The number of stem cells in the vegetative meristem of the L2 cell layer, which ultimately contribute to the germline, is estimated to be <10 cells (Yu et al., 2024), and these cells undergo ∼10 mitoses between branching (Burian et al., 2016). This population of stem cells must eventually expand via mitosis during flower development and finally undergo meiosis to generate the pollen and egg cells. The number of pollen (2000-8000) per plant provides an order of magnitude estimate for the cell population size of the germline (Tsuchimatsu et al., 2020). Our simulation resulted in 1-2 mutations per genome/generation, similar to empirical measures from Arabidopsis mutation accumulation experiments.

### Selection and mutation bias affect genic mutation rates

As expected, modeling lower genic mutation rates resulted in lower observed mutation rates in genic regions, but not in intergenic regions (**Figure 1b**,**c**). Selection on non-synonymous mutations was inconsequential in cases where the mean selection coefficient (*s*) is smaller than 0.03, which is consistent with the small effective population size of meristem cells destined for the germline during most of development and the extreme population bottleneck (Ne = 1, single seed descent) imposed in each generation. Empirical estimates of the distribution of fitness effects of deleterious mutations provide realistic bounds for the mean selection coefficient *(s)* on the order of −0.0001 to −0.01. It is therefore unlikely that selection would have a strong effect on mutation rate estimates (see **Table S2**) (Böndel et al., 2019; Eyre-Walker et al., 2006; Kibota & Lynch, 1996; Kim et al., 2017; Perfeito et al., 2014; Plavskin et al., 2024; Robert et al., 2018; Saber et al., 2025a; Williamson et al., 2014).

The effect of selection is also dependent on the degree of dominance of deleterious alleles. The effect of selection on strongly deleterious alleles (s > 0.03) is still weak when deleterious alleles are recessive (dominance, D = 0). This is expected since during development, mutations arising in somatic cells are necessarily heterozygous. The largest surveys of somatic mutation and evolution come from the field of cancer biology, which find that negative selection is only detectable in hemizygous genes, consistent with the recessive nature of most deleterious mutations (Bakhoum & Landau, 2017; Martincorena et al., 2017; Van den Eynden et al., 2016). Such hemizygosity would not exist in inbred Arabidopsis plants used for mutation accumulation. Thus, developmental selection is expected to be weak in Arabidopsis due to the combination of small effective population sizes of initial cells in the meristem, as well as the strength of selection against the recessive nature of deleterious mutations.

Nevertheless, selection can have consequences on observed gene body mutation rates under extreme conditions; when mean selection coefficients are extremely strong and dominant, observed mutation rates in gene bodies were lower than the neutral expectation determined by the parameters set in the simulation (**Figure 1d**). These conditions provided an opportunity to test whether such selection could be detected and accounted for to accurately estimate true mutation biases (degree of genic hypomutation independent of selection). When non-synonymous sites were removed by selection, we detected significant deviations from the neutral NS/S (**Figure 1e-f**). By accounting for this selection, as described above, we can accurately estimate the true degree of genic hypomutation (**Figure 1g**).

We also found that individual classes of variants could help distinguish selection and hypomutation. As expected, under conditions of extreme negative selection with dominant mutations, the loss of non-synonymous mutations surpassed the expected degree of hypomutation due to the mutation rate alone (**Figure 2a**). In contrast, the reduction of synonymous and non-coding mutations matched the true genic hypomutation parameterized in the simulations (**Figure 2b-c**).

**Figure 2.**
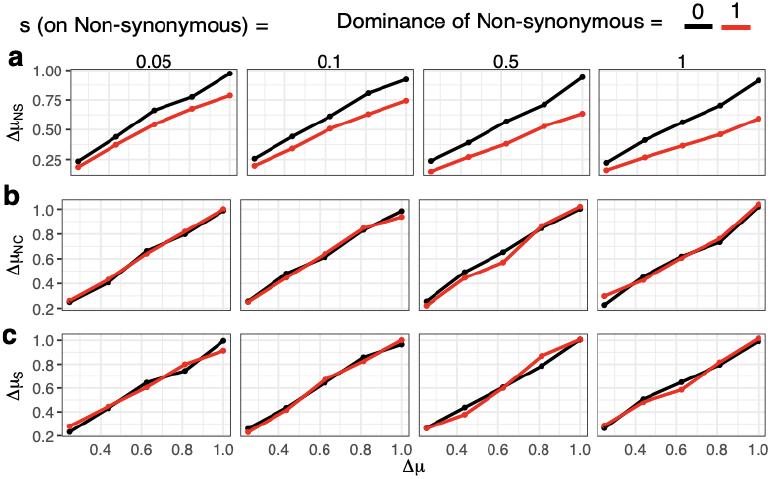
Measuring mutation rate reduction on individual components of genic regions. Visualization only includes simulations with strongly deleterious non-synonymous mutations (s>0.03) **a)** The reduction of non-synonymous mutations (Δμ_NS_) **b)** The reduction of non-coding genic (i.e. introns or untranslated regions) mutations (Δμ_NC_) **c)** The reduction of synonymous mutations (Δμ_S_).

In summary, negative selection is unlikely to have a significant impact on mutation accumulation experiments in plants, given the small effective population sizes of cells contributing to the germline, realistic estimates of selection coefficients, and the dominance of deleterious alleles. Indeed, studies of somatic mutation in plants fail to show purifying selection (Wang et al., 2024). Still, the effects of purifying selection can be pronounced under extreme parameters. In the face of such conditions, correcting for selection against non-synonymous mutations (Δµ_Est_) and directly estimating hypomutation from synonymous (Δµ_S_) and non-coding genic (Δµ_NC_) mutations provides measures of mutation rate heterogeneity independent of selection against non-synonymous mutations (**Figure 1e, Figure 2b**,**c**).

### Evaluating selection in Arabidopsis mutation accumulation data

Mutation accumulation experiments in *Arabidopsis thaliana* reflect the sum of somatic and gametic selection influencing the mutations ultimately observed. We re-analyzed mutation accumulation datasets from independent studies (**Figure 3**, Table S3), constituting a dataset of 7,836 single-nucleotide substitution mutations (*Materials & Methods*) (Belfield et al., 2018, 2021; Jiang et al., 2014; Lu et al., 2021; Monroe et al., 2022; Weng et al., 2019; Willing et al., 2016; S. Yang et al., 2015; Zhu et al., 2021). From these, we estimated the neutral NS/S mutation ratio using methods similar to both classic tools for calculating Dn/Ds (Z. Yang, 2007b), as well as methods previously described that account for substitution frequencies (Martincorena et al., 2017; Rheinbay et al., 2020; Satake et al., 2023) (Materials and Methods). Mutation probabilities, in relation to spectra (i.e., C>T vs T>C), are projected onto all coding sequences, which are randomly sampled, resulting in either a change in amino acid sequence (non-synonymous) or not (synonymous). We used the mutation spectra of non-coding regions (to preclude any effects of selection in coding regions skewing the neutral mutation spectrum). These mutation spectra were scaled to reflect the nucleotide composition of non-coding regions, to control for differences in nucleotide composition between coding and non-coding regions. For all coding regions, we then permuted mutations based on the probability of every possible mutation and defined whether the resulting mutation would be synonymous or non-synonymous. Consistent with expectations, C>T and G>A mutations were the most frequent class of mutation (**Figure 4a**). Given the redundancies of the amino acid code, T>C and C>T mutations were the least likely to cause non-synonymous mutations, as expected (**Figure 4a**).

**Figure 3.**
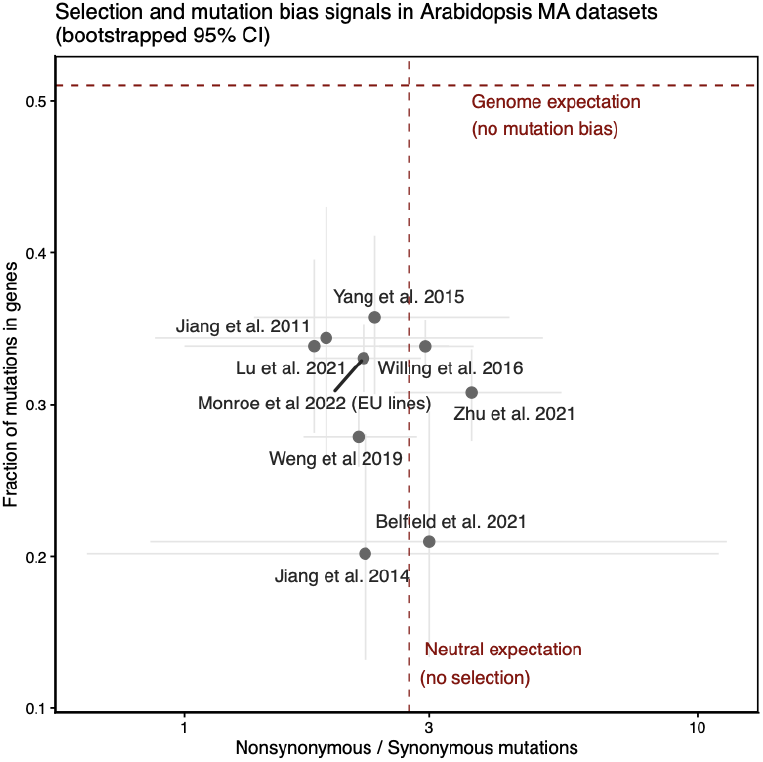
Genic mutation fraction vs. nonsynonymous/synonymous (NS/S) ratio for individual Arabidopsis MA datasets (points with 95% bootstrapped Cis, B = 1,000). The vertical dashed line (NS/S = 2.71) is the neutral expectation from spectrum × codon use; the horizontal dashed line (0.501) is the genome-wide genic fraction. All datasets fall below the genic expectation (consistent with a genic mutation bias), but all NS/S Cis overlap the neutral expectation (i.e. we do not detect systematic purifying selection on coding mutations in any dataset). Genic fraction and NS/S were not correlated across datasets: those with a lower fraction of mutations in genes did not show a correspondingly lower NS/S ratio.

**Figure 4.**
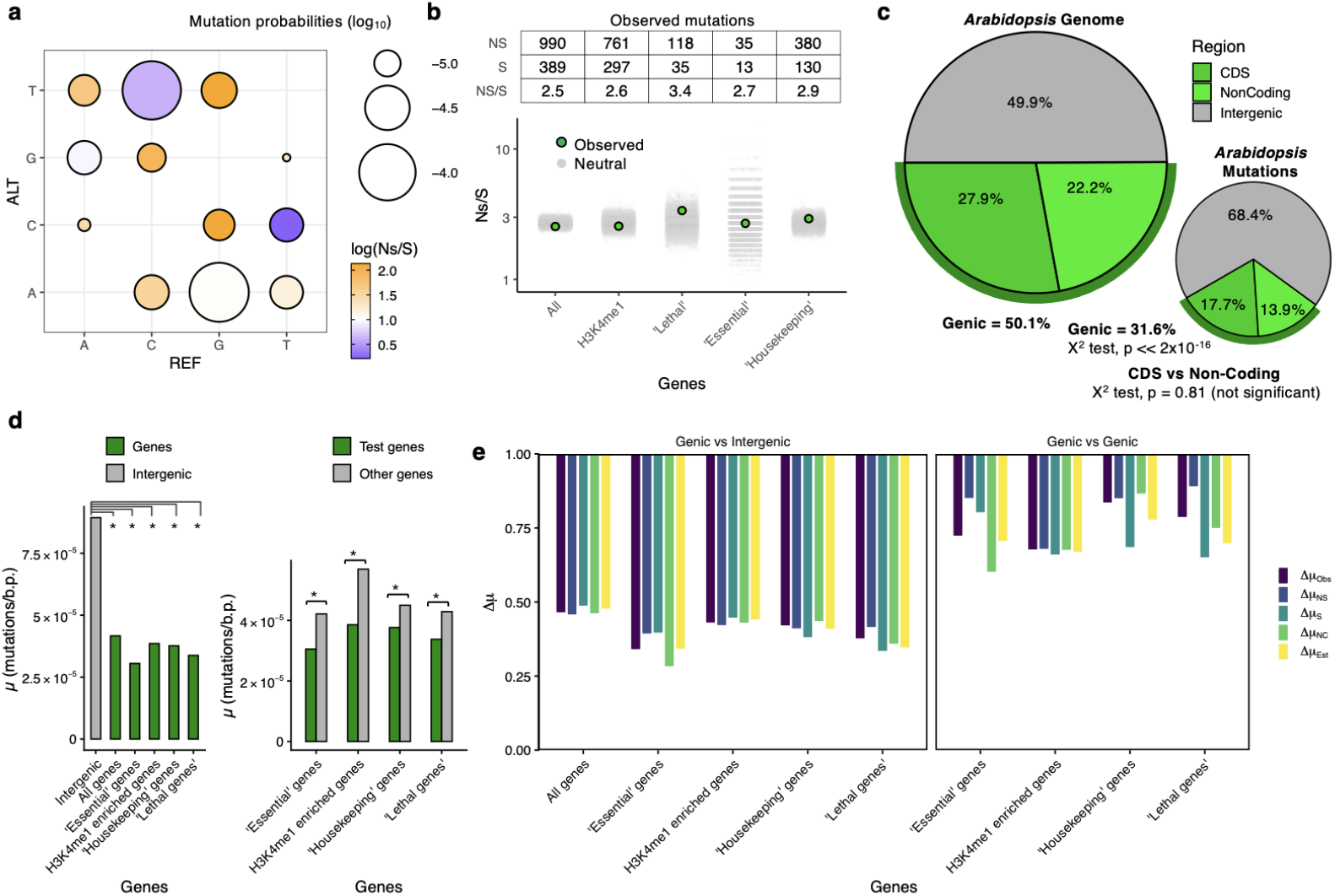
Selection and mutation bias in Arabidopsis mutation datasets. **a)** Projection of mutation spectra (REF > ALT) from non-coding regions onto Arabidopsis coding regions for estimating neutral Ns/S ratios, **b)** Random sampling of coding sequence mutations based on mutation probabilities (1000 iterations), with mutation number samples equal to the number of coding mutations found in all genes (n=1385 mutations), ‘essential’ genes (n=48), ‘lethal’ genes (n=154), H3K4me1-enriched genes (n=1062), and ‘housekeeping’ genes (n=510). **c)** Genome of Arabidopsis (TAIRI0) (upper pie chart) and the proportion of single base substitution (SBS) mutations in each region (lower pie chart). Mutation datasets are from Weng et al. 2019, Belfield et al. 2021, etc. Total mutations, n=7836. **d)** Same gene groups as in (b), comparing mutation rates in gene body (CDS + Non-Coding) with intergenic regions. The left panel shows the comparison of mutation rates in specific gene groups with other genes (i.e. ‘Essential’ genes vs non-essential genes). Asterisks indicate p-value < 0.001 for chi-squared test, **e)** Measures of mutation rate reduction of individual components. For genic vs intergenic comparisons, the mean of sampled neutral Ns/S for all genes from (c) was used (left). The right panel shows genic comparisons of individual gene groups, with neutral Ns/S in these cases indicating the Ns/S observed in other genes for each comparison.

To estimate expected NS/S under neutrality, we randomly sampled (bootstrap n = 1000) synthetic “mutations” in coding sequences, matching the total number of coding region mutations observed in the Arabidopsis mutation dataset for all genes and subsets of genes (e.g., mutations in ‘Essential’ genes only). For all genes, as well as gene subsets, the observed NS/S ratio was within the 95% bootstrapping confidence interval for neutrality. The NS/S of mutation accumulation datasets in Arabidopsis do not show evidence of being subject to significant purifying selection, which is consistent with the expectation for weak efficacy of selection in these experiments. The observed Ns/S ratio in essential and related genes was not lower than the genome-wide average, which is opposite to the prediction made by the developmental selection hypothesis (Majic & Payne, 2023).

#### Mutation rates and selection in gene bodies and functional genes

While gene bodies (coding regions, untranslated exons, and introns) constitute 51.1% of the Arabidopsis genome, they contributed only 31.6% of mutations (**Figure 4c**). This reduction was significant for both CDS (χ^2^ = 686, p < 2×10^−16^) and non-coding (χ^2^ = 581, p-value < 2×10^−16^) genic regions, and was not different between CDS and non-coding genic regions (χ^2^ = 0.06, p = 0.8131)(**Figure 4c**). Thus, coding and non-coding genic features showed the same reduction of mutation rates, consistent with mechanisms potentially contributing to genic hypomutation, such as transcription-coupled repair (TCR) and H3K4me1-recruited DNA repair. Notably, H3K4me1 does not distinguish exons and introns (Oya et al., 2022; Quiroz et al., 2024; Zhang et al., 2009) (Oya et al., 2022; Quiroz et al., 2024; Zhang et al., 2009). The reduction of mutation rates in both exons and introns in plants is distinct from humans, where H3K36me3 specifically marks exons and recruits DNA repair, lowering mutation rates in exons to a greater extent (Frigola et al., 2017; Huang et al., 2018; Li et al., 2013; Moore et al., 2021). Consistent with work previously reported, we observed a ∼53% reduction in genic mutation relative to that observed in intergenic regions (Δµ_OBS_ = 0.53) across all genes (Monroe et al., 2022, 2023; Quiroz et al., 2024). That this reduction was observed in both coding and non-coding genic elements, along with the neutral NS/S of genes, suggests a reduction in mutation rates rather than the effects of selection (**Figure 4b-e**). This is consistent with mutation rate reductions measured directly in soma, which show lower mutation rates in gene bodies, across both exons and introns (Amundson et al., 2023; Davis et al., 2025; Meyer et al., 2025). The expected neutrality of NS/S for all genes was 2.71 (**Figure 3, 4b**). Therefore, reducing gene body mutations by more than 50% via purifying selection would require the removal of nearly all non-synonymous mutations, which is inconsistent with the data.

We also examined the mutation rates of gene bodies for specific sets of genes. H3K4me1 is recognized by DNA repair in plants and is enriched in essential genes (“Essential”) (J. Lloyd & Meinke, 2012), genes predicted to have lethal effects if knocked out (“Lethal”)(J. P. Lloyd et al., 2015), and constitutively expressed (“Housekeeping”; expression detected in 100% of tissues sampled) genes (Mergner et al., 2020). For all of these individual subsets of genes, we observed lower mutation rates than intergenic regions (X^2^ test, p<0.001, **Figure 4d-e**). We also compared the gene body mutation rates of each of these gene sets with those of the rest of the genes in the genome. For all cases, we observed significantly lower gene body mutation rates in each gene set (X^2^ test, p<0.001, **Figure 4d-e**), with observed hypomutation (Δµ_OBS_) ranging from ∼0.15 to ∼0.3.

We then tested and found no evidence that these reductions were due to selection. This is apparent in the observation that these gene sets have a higher Ns/S ratio than the genome-wide average, which is the opposite of awhat would be expected if they were under stronger purifying selection (**Figure 4b**). We also measured the individual components of hypomutation. For these comparisons of mutation rates between genes, we used the observed Ns/S in other genes as the neutral expectation. For example, in comparing essential versus non-essential genes, the N/S of non-essential genes (2.55) was used as the neutral expectation. We found that for all cases, mutation rates were lower regardless of measuring non-synonymous (Δµ_NS_), synonymous (Δµ_S_), or non-coding genic (Δµ_NC_) mutations specifically (**Figure 4e**). This was true for all genes and subsets when comparing against intergenic regions, as well as for individual gene sets when comparing between genes (**Figure 4e**). Given the apparent neutrality of mutations in these genes, the mutation reduction estimate (Δµ_Obs_) was comparable to that calculated after accounting for selection (Δµ_Est_), as expected.

## Discussion

Our framework models selection through comparisons of non-synonymous and synonymous mutations. While this is a practical and widely applied approach (Eyre-Walker & Keightley, 2007; Kimura, 1977; Lawson et al., 2024; Satake et al., 2023; Z. Yang, 2007b), it ignores selection acting on synonymous, intronic, and regulatory mutations. Empirical estimates indicate that selection on synonymous mutations is much weaker than that on non-synonymous mutations (Böndel et al., 2019; Z. Chen et al., 2017; Keightley et al., 2005; Kim et al., 2017). Testing NS/S against neutral expectations helps to detect selection where it occurs, but also provides an upper bound. If non-synonymous sites appear to accumulate neutrally, weaker targets are expected to certainly do so as well.

These developmental simulations improve upon prior models (Majic & Payne, 2023), 2023) by incorporating realistic meristem population sizes; however, they remain simplifications. In Arabidopsis, each branch or floral meristem arises from a single L2-layer stem cell, creating frequent and severe bottlenecks (Burian et al., 2016; Y. Chen et al., 2024; He et al., 2024; Johannes, 2025a, 2025c; Reusch et al., 2021)). These events likely increase the role of drift while reducing the efficacy of somatic selection, although the stochastic choice of founder cells at branch formation could still allow for limited lineage selection. Some degree of competition among branches for resources might also generate selective pressures at the whole-organism level that are not captured in cell-based simulations. More detailed models incorporating such branching hierarchies and resource dynamics would better reflect development processes and selection, especially in larger, longer lived plants.

Finally, our analysis focuses on single-base substitutions, but other mutational processes—such as transposable element (TE) insertions and structural variants—also exhibit strong, chromatin-dependent biases. TE families often target intergenic, promoter-proximal, or genic regions depending on chromatin context (Davis et al., 2025; Huang et al., 2022; Movilli et al., 2025; Quadrana et al., 2016, 2019). Likewise, small indel rates are influenced by local sequence and chromatin structure. Expanding correction frameworks to include these mutation classes will be key to building a more comprehensive model of how DNA repair and selection shape mutational landscapes in plants.

Discoveries of lower mutation rates in active gene bodies are consistent with mechanisms of transcription-coupled and epigenome-recruited repair (Martincorena & Luscombe, 2013). In plants, genes are protected from mutation by DNA repair proteins that preferentially repair transcribed genes (Oztas et al., 2018) and complexes that bind H3K4me1 via Tudor histone reader domains (Belfield et al., 2018; Quiroz et al., 2024). Importantly, “Essential”, “Lethal”, and “Housekeeping” genes are all enriched for H3K4me1 (Monroe et al., 2022). Transcription-coupled repair provides an additional mechanism that reduces mutation rates in genic regions, regardless of the consequences (non-synonymous, synonymous, or non-coding) (Oztas et al., 2018).

The histone variant H2A.X is also enriched in euchromatin and gene bodies, which, when phosphorylated to γ-H2A.X, is bound by BRCT-domain proteins that promote break repair (Amiard et al., 2010; Vegesna et al., 2025). Similarly, acetylated H4, H2A, and H2A.Z histone modifications enriched in euchromatic, gene-rich regions are bound by YEATS-domain readers, which promote efficient homologous recombination repair (Vegesna et al., 2025; Yelagandula et al., 2014). Thus, mechanisms that enhance DNA repair in transcriptionally active regions offer experimental evidence for genic hypomutation beyond post hoc selection, whether during development or across generations, in plant mutation accumulation studies.

Developmental selection within meristems is expected to be weak for theoretical and empirical reasons. The effective population size of shoot apical meristem stem cells is extremely small—often fewer than ten cells—so the efficacy of selection is minimal relative to drift (Burian et al., 2016; Lanfear, 2018). Cell fate is largely governed by positional cues and hormone gradients within rigid tissues constrained by cell walls, which prevents spatial competition among daughter cells (Kitagawa & Jackson, 2019; Meyerowitz, 1997). Although dominant lethal mutations could eliminate individual cells, such events are rare, and most deleterious alleles are recessive with selection coefficients typically <0.01 (Table S1), The tendency of more deleterious mutations to exhibit greater recessivity than less deleterious mutations is evident in *Arabiopsis*, as well as other taxa such as *Drosophila* and *C. elegans* (Caballero & Keightley, 1994; Halligan & Keightley, 2009; Huber et al., 2018; Saber et al., 2025b; Schultz et al., 1999). Consistent with this, somatic mutation studies in humans and clonally propagated plants reveal little to no evidence of purifying selection, but do show signals of positive selection (Beaulieu et al., 2025; Lawson et al., 2025; Martincorena et al., 2015; Meyer et al., 2025; Quiroz et al., 2024). Together, these findings suggest that developmental selection within plant meristems is probably too weak to explain widespread genic hypomutation, reinforcing DNA repair as the more parsimonious mechanism.

Another developmental context where selection could act on new mutations is the haploid gametic phase, which can directly expose recessive deleterious mutations to selection, influencing pollen viability and success. In *Arabidopsis*, however, self-compatibility and high levels of homozygosity greatly reduce pollen competition by nature of having a reduced number of genotypes, resulting in weaker selection on pollen-expressed genes, which also tend to exhibit higher tissue specificity and relaxed purifying selection compared to sporophytic genes. Consequently, while haploid selection could potentially expose recessive deleterious alleles, its effects in *A. thaliana* are expected to be minimal and unlikely to alter the observed patterns of genic hypomutation (D. Charlesworth & Charlesworth, 1992; Harrison et al., 2019).

Even with strong evidence for preferential DNA repair in transcribed and euchromatic regions, and no indication that purifying selection explains genic hypomutation, it remains important to formally account for potential selective effects. Because non-synonymous mutations constitute the principal targets of purifying selection (Kimura, 1977; Z. Yang, 2007a), they provide a natural basis for correction. Our forward simulations demonstrate that adjusting for the potential removal of non-synonymous variants effectively eliminates spurious biases without distorting true mutation rate differences. In *Arabidopsis thaliana*, where mutation accumulation experiments consistently show no detectable selection on either coding or noncoding mutations, this correction has negligible influence on the estimated degree of genic hypomutation—further confirming that lower mutation rates in genes primarily reflect mutational (e.g., damage, DNA repair) processes rather than selection. Nevertheless, quantitative methods incorporating such corrections remain valuable in systems with larger effective population sizes or stronger selection, where purifying selection could meaningfully shape observed mutation spectra.

In natural populations, background selection can complicate the interpretation of differences in genic mutation rates. By removing linked neutral variants near deleterious alleles, especially in regions of low recombination (B. Charlesworth et al., 1993; Nordborg et al., 1996), background selection can reduce both non-synonymous and synonymous polymorphisms. However, without a very high recombination frequency, these effects can extend broadly across the genome, blurring fine-scale distinction between neighboring genic and intergenic regions at the level of polymorphism data. Mutation accumulation lines maintained through strict inbreeding and minimal recombination are largely insulated from background selection. As a result, they provide a clearer view of true mutation rate heterogeneity, free from the confounding effects that could obscure such patterns in natural populations.

In conclusion, mutation biases such as genic hypomutation can shape the origins of genetic diversity. Here, we present a simple quantitative framework to distinguish between intrinsic mutation bias and purifying selection on coding sequences, and to evaluate the potential for selection to act within the plant soma. Future work to study how DNA damage, repair, and development together define where and how new variation arises is essential toward understanding the evolution of plant genomes.

## Materials and Methods

### Simulating selection and mutation rate heterogeneity in Arabidopsis mutation accumulation lines

To model the accumulation of mutations in Arabidopsis thaliana, we utilized SLiM version 4.1 for forward evolutionary simulations. SLiM enables us to simulate evolutionary scenarios that capture developmental selection with parameters to quantify the effects on mutation accumulation.

The parameters and settings for our simulations included both mutation and selection heterogeneity. We used a mutation multiplier (M) to adjust the mutation rate in genic regions, with values sampled from 0.25 to 1.0 in increments of 0.1875. The selection coefficients for negative selection were drawn from gamma distributions with different means for non-synonymous mutations (*s*). Specifically, *s* values for non-synonymous mutations were sampled from a range of neutral to lethal {0, 0.001, 0.01, 0.03, 0.05, 0.1, 0.5, 1}. Note: Empirical estimates of mean selection coefficients are on the scale of 0.001 to 0.01. Dominance coefficients (D) were also varied, with D values for non-synonymous mutations sampled from {0, 0.1, 0.5, 1}. The shape parameter of the gamma distribution was set to 0.14, based on previous work (Plavskin et al.,(Plavskin et al., 2024). Simulated genomes were 8,000 base pairs (bp) in size, with 50% of the region designated as genic regions. Of these genic regions, 50% were coding, and within coding regions, 2/3 of mutations were non-synonymous.

The simulation process began with initializing the nucleotide-based settings and an ancestral genome of 8,000 bp. We defined mutation types for different regions: neutral mutations for intergenic regions, deleterious mutations for non-synonymous regions, and similarly specified parameters for synonymous and non-coding genic regions. Genomic elements were assigned specific mutation rates and divided into non-genic (the first 4000 bp) and genic (the last 4000 bp) regions. Intergenic mutation rates were set to 3.5×10^−6^, which yielded ∼1-2 mutations/genome/generation, which is similar to experimentally reported mutation rates in Arabidopsis (Weng et al., 2019).

We simulated population dynamics, starting with 2 cells, which represent the genetically effective cells in an embryo at generation 1 (G1). The cell population expanded through several stages, mimicking the development from the central zone of the meristem to gametogenic cells. At specific time steps, cells underwent sexual reproduction, simulating zygote formation and subsequent embryonic development. Each simulation represented three generations of mutation accumulation. Population sizes and cloning rates were adjusted at specific intervals to accurately reflect biological processes.

After three generations, we sampled the population and recorded fixed mutations. Each simulation was run 4000 times for every combination of parameters (S, D, M).

For each parameter combination, we quantified mutation rates in genic and intergenic regions and recorded the number of observed non-synonymous and synonymous mutations. These values were compared against the neutral expectation to assess deviations attributable to selection using chi-squared tests. We then measured the observed reduction in mutation rates within gene bodies—both overall and separately for non-synonymous, synonymous, and non-coding genic sites—and estimated the true degree of genic hypomutation after correcting for the selective loss of non-synonymous mutations. Finally, we compared these corrected estimates with the known true mutation rates specified in the simulations to evaluate the accuracy and performance of our correction framework.

### Testing for selection and mutation bias in Arabidopsis thaliana mutation accumulation experiments

We compiled and re-analyzed single-nucleotide substitution (SBS) data from nine published Arabidopsis thaliana mutation accumulation (MA) experiments, each aligned to the TAIR10 reference genome. (Belfield et al., 2018, 2021; Jiang et al., 2014; Lu et al., 2021; Monroe et al., 2022; Ossowski et al., 2010; Weng et al., 2019; Willing et al., 2016; S. Yang et al., 2015; Zhu et al., 2021). To ensure accuracy, we removed duplicate mutation calls identified across studies, which may represent shared sequencing or annotation artifacts. After filtering, a total of 7,836 unique SBSs remained for analysis. Structural variants (SVs) and indels were excluded, as our focus was on base substitutions; potential selection against SVs or indels would therefore not affect our results. SBS from published mutation-accumulation or resequencing studies: Belfield et al. 2021 (n = 105), Jiang et al. 2011 (n = 128), Jiang et al. 2014(n = 114), Lu et al. 2021 (n = 263), Monroe et al. 2022, EU lines (n = 1,670), Weng et al. 2019 (n = 1,865), Willing et al. 2016 (n = 2,495), Yang et al. 2015 (n = 316), and Zhu et al. 2021 (n = 880).

We annotated all retained single-base substitutions (SBS) in relation to the *Arabidopsis thaliana* TAIR10 reference genome assembly and annotation. Each mutation was classified as genic or intergenic, and genic sites were further partitioned into coding sequence (CDS) and genic non-coding sequence (introns and UTRs). Mutations were also intersected with biologically defined gene sets from the literature, including essential genes (J. Lloyd & Meinke, 2012), lethal genes (Lloyd et al. 205), H3K4me1-enriched genes (Niu et al. 2021), and broadly expressed/housekeeping genes profiled across 54 tissues (Mergner et al. 2016). For mutations that mapped to CDS, we reconstructed the corresponding TAIR10 coding sequence *in silico*, introduced the observed nucleotide substitution, retranslated the modified CDS, and classified the effect as nonsynonymous (amino-acid altering) or synonymous. To obtain a neutral, sequence-aware expectation for the nonsynonymous/synonymous (NS/S) ratio, we combined two sources of information: (i) the empirical *A. thaliana* SBS spectrum estimated from the compiled mutation datasets, and (ii) genome-wide TAIR10 codon usage. We simulated mutations *in silico* by sampling substitutions probabilistically according to the empirical spectrum and placing them onto the TAIR10 codons, determining for each simulated event whether it would produce a synonymous or nonsynonymous change after translation. This was repeated 10,000 times for each focal gene set (all genes, essential, lethal, H3K4me1-enriched, broadly expressed), always matching the number of observed coding mutations in that set. The resulting bootstrap-like distributions provided neutral NS/S expectations and confidence intervals that explicitly account for both mutation biases (which bases mutate to what) and the codon composition of the genes being tested. We then compared the observed NS/S of each gene set to its corresponding neutral distribution and recorded the proportion of neutral draws as or more extreme than observed as an empirical p-like measure of depletion/enrichment.

After annotation and neutral benchmarking, we applied the selection-aware hypomutation framework described in the main text to every focal gene set. For each set, we first computed an observed genic mutation rate (observed mutations divided by total genic bp for that set) and compared it to a reference mutation rate. For gene-to-gene comparisons, the reference was “all other genes”; for gene-to-intergenic comparisons, the reference was the intergenic sequence. This yielded the observed genic-to-reference multiplier (Δµ_o_ < 1 implies hypomutation). We then asked whether Δµ_o_ could be explained purely by the loss of nonsynonymous mutations to selection. Using (i) the neutral NS/S estimate (either from the non-focal genes or from the in silico permutations) and (ii) the fraction of genic sequence that is coding in the focal set, we calculated how many nonsynonymous, synonymous, and non-coding genic mutations should have been observed if the focal set had mutated at the reference rate. The shortfall in nonsynonymous mutations was converted to a selection-correction term and added back to Δµ_o_ to obtain the selection-corrected genic hypomutation estimate (ΔµEst). In parallel, we applied the same logic to the individual mutation classes to obtain component estimates ΔµNS, ΔµS, and ΔµNC, which allowed us to test whether hypomutation persists in synonymous or non-coding genic sites where selection is expected to be weak. For genome-wide genic vs. intergenic contrasts, we used the mean NS/S across genes as the neutral baseline; for pairwise tests between gene subsets, we used the comparison group’s NS/S as the neutral baseline.

## Acknowledgments

The Monroe Lab is supported by National Science Foundation grants 2317191 and 2338236 to J.G.M.. Research was conducted at the University of California Davis, which is located on land that was the home of the Patwin people for thousands of years.

## Data and Code availability

Code for this study is available on GitHub at github.com/greymonroe/At_devel_sel.

## Supplemental tables

**Supplementary Table 1**. Results from SLiM simulations of somatic meristem selection and mutation accumulation

**Supplementary Table 2**. Empirical estimates of distribution of fitness effects, expected strength of deleterious selection (s).

**Supplementary Table 3**. Mutation call-set (SBS) from *Arabidopsis thaliana* experiments

**Supplementary Table 4**.Mutation probabilities, effects, based on codon use and mutation spectra

